# Evidence of a dysregulated Vitamin D pathway in SARS-CoV-2 infected patient’s lung cells

**DOI:** 10.1101/2020.12.21.423733

**Authors:** Bijesh George, Ravikumar Amjesh, Aswathy Mary Paul, T. R. Santhosh Kumar, Madhavan Radhakrishna Pillai, Rakesh Kumar

## Abstract

Although a defective vitamin D pathway has been widely suspected to be associated in SARS-CoV-2 pathobiology, the status of the vitamin D pathway and vitamin D-modulated genes in lung cells of patients infected with SARS-CoV-2 remains unknown. To understand the significance of the vitamin D pathway in SARS-CoV-2 pathobiology, computational approaches were applied to transcriptomic datasets from bronchoalveolar lavage fluid (BALF) cells of such patients or healthy individuals. Levels of vitamin D receptor, retinoid X receptor, and CYP27A1 in BALF cells of patients infected with SARS-CoV-2 were found to be reduced. Additionally, 107 differentially expressed, predominantly downregulated genes modulated by vitamin D were identified in transcriptomic datasets from patient’s cells. Further analysis of differentially expressed genes provided eight novel genes with a conserved motif with vitamin D-responsive elements, implying the role of both direct and indirect mechanisms of gene expression by the dysregulated vitamin D pathway in SARS-CoV-2-infected cells. Network analysis of differentially expressed vitamin D-modulated genes identified pathways in the immune system, NF-κB/cytokine signaling, and cell cycle regulation as top predicted pathways that might be affected in the cells of such patients. In brief, the results provided computational evidence to implicate a dysregulated vitamin D pathway in the pathobiology of SARS-CoV-2 infection.

## Introduction

Covid-19 (SARS-CoV-2) pandemic continues to be a major global public health crisis and has caused 1,605,091 mortalities as of January 9, 2021^1^. Scientifically validated medicine and/or vaccine that might be suitable for the society at-large are about to be made available. Simultaneously, the scientific community is being largely guided by the epidemiological, symptomatic, socioeconomic, and population variabilities, and predictive disease models for understanding the cellular basis of infectivity and emerging patterns of the pandemic. This includes the studies of reinfection and/or coinfection with other viruses and formulation of counter measures to minimize the probability of SARS-CoV-2 infection^2–4^. Several recent studies have provided clues about the nature of cellular pathways that might be responsible for the evident population variability in terms of susceptibility, exhibition of symptoms, severity, progression of the disease, and recovery from the infection^5,6^. This process is further benefitted by applying the lessons gained from the interactions between the human cells and other single-stranded RNA viruses in hosts such as those causing influenza, hepatitis, Ebola, and AIDS.

One of the correlations that has recently gained scientific attention is the correlation between vitamin D deficiency and SARS-CoV-2 infection. This is partly because of the role of vitamin D in cellular phenotypes, including immunity modification and inflammation, and its ability to influence the endothelial cell biology and the functionality of ACE2, one of the major receptors for SARS-CoV-2^7–9^. Although correlative studies between vitamin D pathway and SARS-CoV-2 continue to provide positive clues about the significance of vitamin D in SARS-CoV-2 pathobiology, the expression status of core components of the vitamin D pathway, and hence, the functionality of the pathway in patients infected with SARS-CoV-2 remain unknown.

Vitamin D is one of the most studied steroid hormones with various cellular functions. The precursor vitamin D3 is generated from 7-hydrocholestetol by a nonenzymatic reaction catalyzed by ultraviolet B radiation. In the liver, vitamin D3 is modified to 25-hydroxy vitamin D and further to 1,25-dihydroxyvitamin D (calcitriol) in kidneys as well as in other organ systems in the presence of CYP27B1 enzyme. Cellular effects of vitamin D result from its nongenomic or cytoplasmic effects and/or genomic nuclear effects. They are caused by 1,25-dihydroxy vitamin D binding to the vitamin D receptor (VDR) or retinoid X receptor (RXR) complex within specific responsive DNA sequences in the target genes or through indirect mechanisms^9^. In addition to the status of vitamin D3/cholecalciferol, biosynthesis of 1,25-dihydroxy vitamin D is positively and negatively modulated by alpha-hydroxylases CYP27B1 and CYP24A1, respectively. Interestingly, these enzymes are under the control of fibroblast growth factor 23 (FGF23), a paracrine growth factor expressed in the lung, heart, and kidney^10^. Increased systemic levels of FGF23 are generally associated with general inflammation, cardiac hypertrophy, chronic kidney disease, and inflammation in lung airway^11^; all these phenotypes are found to be closely associated in SARS-CoV-2 pathogenesis. In general, FGF23 acts through its receptors, FGFR1–4, in its coreceptor klotho-dependent or -independent manner. Other activities of FGF23 relevant to SARS-CoV-2 are as follows: participation in cardiac myocytes in stimulating the levels of fibrotic factors, cardiac fibrosis, and pathological cardiac remodeling^12^; induction of proinflammatory cytokine IL-1beta^11^; inhibition of ACE2 expression, angiotensin II (Ang II) induction, and blood pressure in kidney^13^; and inhibition of M1 to M2 transitioning of macrophage and counteracting anti-inflammation response^14^. Additionally, FGF23 and CYP24A1 are components of a regulatory vitamin D feedback loop because both molecules are genomic targets of 1,25-dihydroxy vitamin D and are stimulated by it in a VDR-dependent manner^11,15,16^. In addition to vitamin D, FGF23 levels have been shown to be induced by obese growth factor leptin^16^, master regulator of hypoxia HIF1-alpha,^17^ and proinflammatory TNF-alpha^17^. For example, leptin induces FGF23 expression and inhibits CYP27B1 expression^18^, implying a potential compromised status of 1,25-hydroxy vitamin D under conditions of obesity. Recent data suggest that increased levels of inflammatory cytokines in SARS-CoV2 infected patients with a reduced level of vitamin D^19^. Currently, we do not know the status of FGF23 and other core modifiers of the vitamin D pathway in lung cells of patients infected with SARS-CoV-2.

For productive infection, many viruses use the host machinery to antagonize the host defense mechanism while using survival pathways to abort apoptosis for ensuring a successful completion of their replication and generation of progeny virions. In this context, previous work suggest that during the host–RNA-virus interaction, pathogen replication and its propagation are profoundly influenced by human p21-activated kinases (PAKs)^20^, which are the established modifiers of cytoskeleton remodeling, inflammation, thrombosis, cell survival, and gene expression or repression, in addition to its oncogenic role in cancer development progression^21–25^. However, to the best of our knowledge, no published data exist about the effect of SARS-CoV-2 on PAK mRNA expression levels in the context of the vitamin D pathway. Current data suggest a mechanistic role of PAK-dependent actin polymerization in vitamin D-mediated stimulation of FGF23 expression^26^; vitamin D signaling stimulates actin depolymerization in endometrial cancer by inhibiting RAC1 and PAK1 expression^27^; the vitamin D pathway uses PAK1 in protecting murine fibroblasts from vitamin E-succinate-triggered apoptosis; and dependency of cholecalciferol-mediated NF-kB transactivation on PAK1 activity^28^. These examples of regulatory crosstalk between vitamin D and PAKs suggest that the cellular effects of vitamin D in the context of SARS-CoV-2 pathobiology might not be merely affected by the biosynthesis of cholecalciferol but also by the status of FGF23, vitamin D metabolic enzymes, and PAK signaling. However, the status of vitamin D pathways in SARS-CoV-2 remains unknown.

To better understand the significance of vitamin D in SARS-CoV-2 pathobiology, we applied computational approaches to study the expression levels of core components and modifiers of the vitamin D pathway and to understand its potential relationship with the status of the PAK pathway in SARS-CoV-2-infected individuals.

## RESULTS

### Status of the vitamin D pathway in viral infection model systems

To better understand the significance of the vitamin D pathway in the pathobiology of SARS-CoV-2 infection, we first examined the expression levels of core components of the vitamin D pathway in various models of viral infection using the Signaling Pathways Project Datasets (SPPD), a collection of transcriptomic datasets initially biocurated by the Nuclear Receptor Signaling Atlas Organization^29^. As illustrated in Fig. 1, we found evidence of both down- and upregulation of molecules belonging to the vitamin D pathway. In general, we noticed a preferential downregulation of the vitamin D pathway in majority of the studies compiled in SPPD. We believe that the noted variability in the levels of molecules among various independent biological studies is a common observation and might arise because of differences in the nature and preparation of test samples as well as experimental conditions and reagents used in the different studies. Specifically, we observed reduced expression of vitamin D coreceptors RXRA, CYP27A1, CYP24A1, and FGFR1-4 in most of the SARS virus-derived datasets. However, most of the SPPD datasets lacked information of the vitamin D pathway in the SARS-CoV-2 infection model system, except for CYP27B1 upregulation. In general, these observations are consistent with the notion of a general reduced expression of several, but not all, components of the vitamin D pathway caused by viruses.

**Figure 1:**
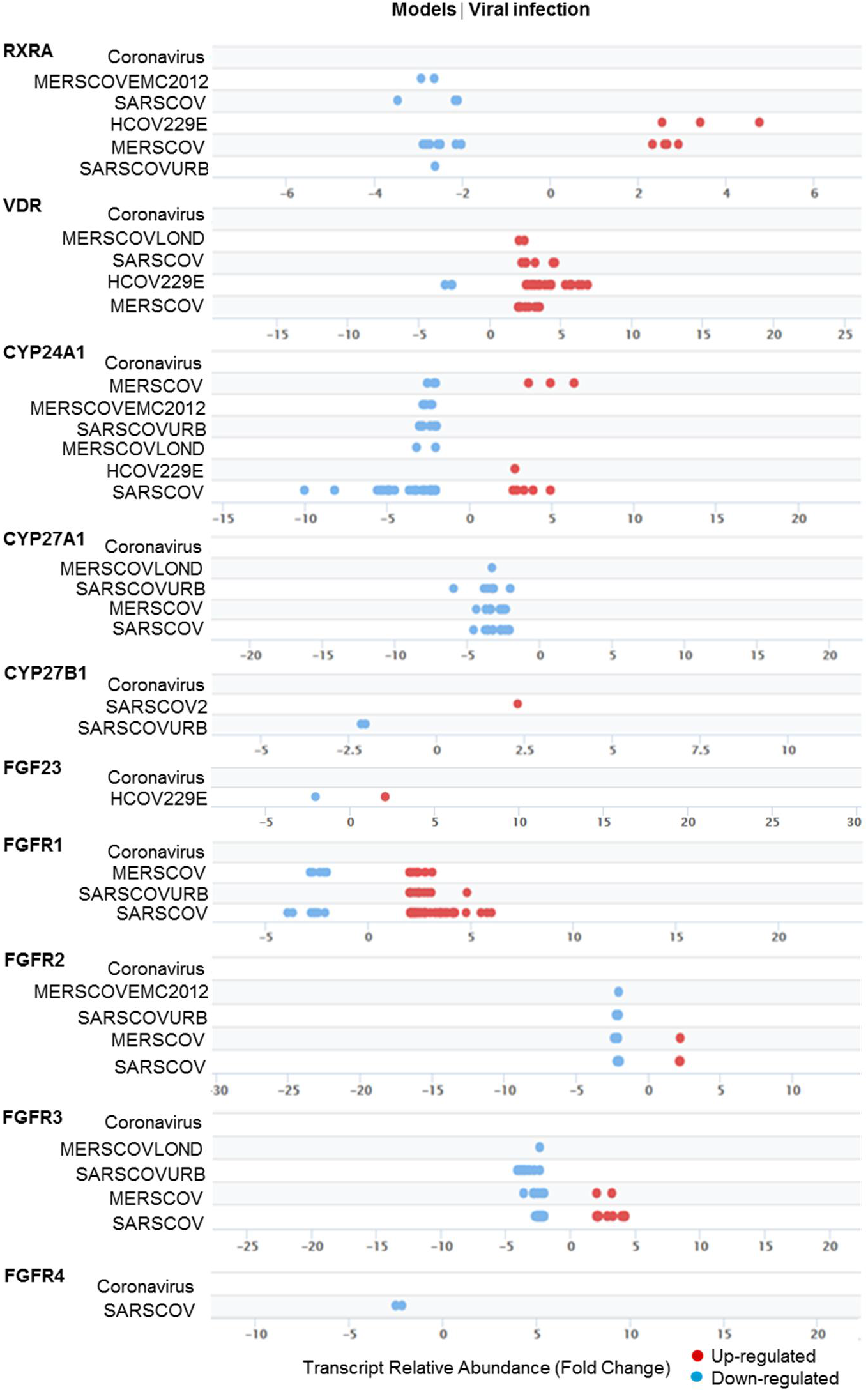
Outlook of Vitamin D components in viral infection models. Expression and distribution of individual genes in Vitamin D component with FDR 5E-02 from vitamin D components under various viral infection models from Signaling Pathway Project.

### Suppression of the vitamin D pathway in lung cells of patients infected with SARS-CoV-2

To assess the status of the vitamin D pathway in lung cells of patients infected with SARS-CoV-2, we next evaluated the levels of core molecules of the vitamin D pathway in three separate RNA sequencing-based transcriptomic datasets of bronchoalveolar lavage fluid (BALF) cells from patients with confirmed SARS-CoV-2 infection and in control healthy subjects^30–32^, as well as in A549, NHBE, and Calu3 human lung cell lines expressing infected SARS-CoV-2^31^. Our analysis indicated an association between SARS-CoV-2 infection and reduced expression of VDR and its mandatory binding partner RXR and reduced expression of CYP27A1 in two of the three human transcriptomic studies and increased expression of CYP24A1 and reduced expression of FGFR1 in only one of the three SARS-CoV-2-infected patients’ transcriptomic studies (Fig. 2a, Supplementary table 1). In contrast to the levels of core components of the vitamin D pathway in BALF cells from SARS-CoV-2-infected patients, infection associated SARS-CoV-2 overexpression in three lung cancer cells did not alter the expression of most components of the vitamin D pathway. However, SARS-CoV-2 overexpression in A549 and Calu3 cells downregulated FGFR1-4 levels (Fig. 2b, Supplementary table 1). These observations suggest that data from cell lines infected with ectopic overexpression by infection of SARS-CoV-2 may not always be compatible with patient-derived primary cultured cells to understand the significance of the vitamin D pathway in SARS-CoV-2-infected patients. Therefore, we focused on transcriptomic data from the patients in subsequent analyses.

**Figure 2:**
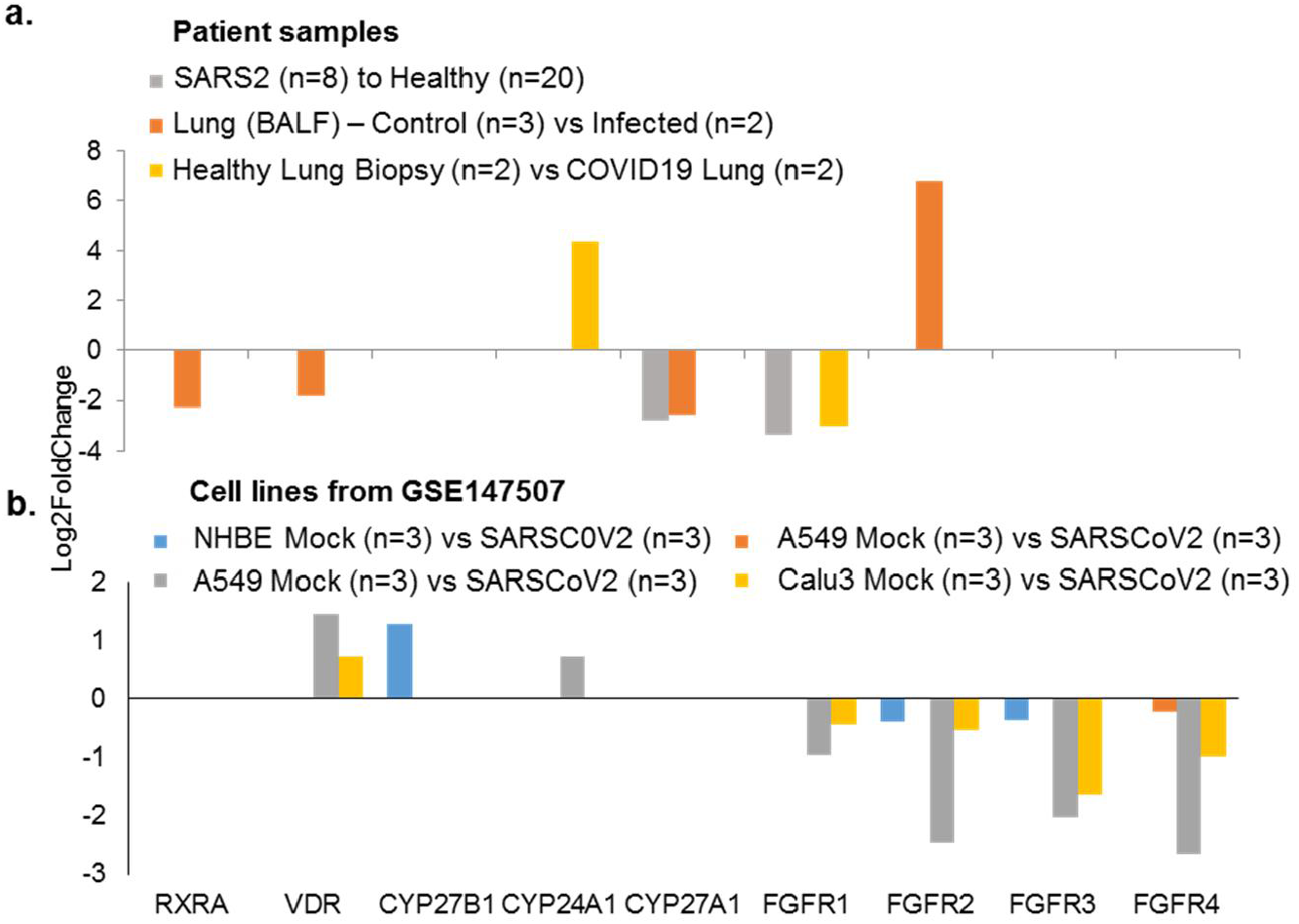
Expression profile of genes under SARS-CoV2 infection. **a)** Bar chart represents the expression profile of Vitamin D components genes from SARS-CoV-2 infected patient samples^30–32^ (PMID – 32407669, PMID – 32228226 and GSE147507) when compared to its healthy controls and **b)**. When the analysis performed in A549, NHBE and Calu3 cell lines from GSE147507.

### Vitamin D-modulated genes overlap with differentially expressed genes in lung cells of patients infected with SARS-CoV-2

Vitamin D mediates its biological effects via upregulating or downregulating the expression of cellular genes responsible for various biological processes. Vitamin D modifies the expression of cellular genes either through a direct mechanism involving predicted or validated VDR motifs in the target genes and/or through indirect pathways^34,35^. Further, we assessed whether noted downregulation of the vitamin D pathway in SARS-CoV-2 might also be accompanied by misregulation of vitamin D-modulated genes, and eventually, resultant functions of such gene products. Thus, as illustrated in Fig. 3a, we found a widespread overlap of vitamin D-modulated genes with differentially expressed genes in three transcriptomic datasets from cells of patients infected with SARS-CoV-2. Upon cross-comparing vitamin D-modulated genes that overlapped in SARS-CoV-2 transcriptomic datasets used here, we recognized 43 differentially expressed vitamin D-modulated genes that shared vitamin D dataset and three transcriptomic datasets from lung cells of patients infected with SARS-CoV-2 (Fig. 3b).

**Figure 3:**
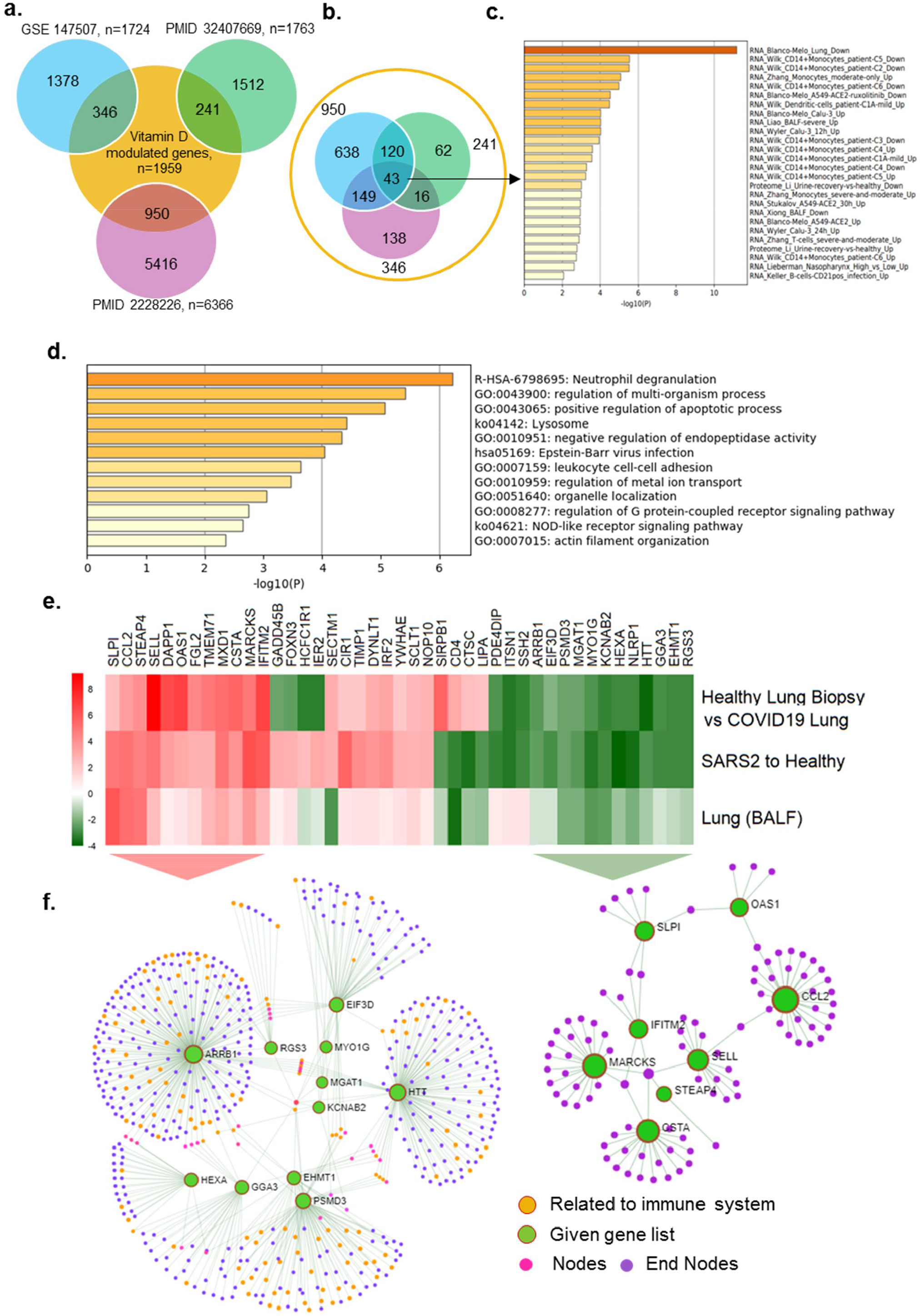
Expression of Vitamin D regulated genes in COVID19 patients. **a)** Number of genes found overlapped with Vitamin D regulated genes and three SARS-COV-2 datasets are denoted. **b)** 43 genes were identified which are common for 3 SARS-COV-2 datasets and also regulated by Vitamin D, **c)** Bar diagram shows the summary of enrichment analysis in SARS-COV-2 from Metascape. **d)** Bar diagram shows the summary of functional enrichment analysis in from Metascape. **e)** The expression status of these 43 genes in SARS-COV-2 patient data is plotted as heat map. **f)** Network analysis of 19 genes found up-regulated in all three patient dataset and also regulated by Vitamin D is given. The functional enrichment analysis of 19 genes using Reactome identified “Activation of the pre-replicative complex” as primary enrichment (rank 1, hits 9/32 pavlue 3.5e-11), Network analysis of 12 genes down-regulated in all three SARS-COV-2 patient transcriptome datasets. Genes are found functionally related to immune system (rank 56, 131/1140 pvalue 1.31e-21) are remarked.

Enrichment analysis of 43 vitamin D-modulated genes in samples collected from COVID patients suggested that many of these genes were downregulated in SARS-CoV-2-infected lung cells^31^ and CD14^+^ monocytes^36^. The associated primary pathways included neutrophil degranulation, regulation of multiorganism processes, and positive regulation of apoptosis^37^ (Fig. 3c, d). They could potentially influence a whole range of functions in SAR-CoV-2-infected lung cells in a vitamin D-sensitive manner (Supplementary tables 2,3). Expression profiling of these 43 genes indicated that 12 vitamin D-modulated genes (HTT, KCNAB2, EHMT1, RGS3, HEXA, NLRP1, GGA3, MYO1G, ARRB1, PSMD3, MGAT1, and EIF3D) were downregulated in 3 SAR-CoV-2 datasets, 9 genes (SIRPB1, CD4, CTSC, LIPA, PDE4DIP, ITSN1, SSH2, IER2, and HCFC1R1) in 2 datasets, and 3 genes (SECTM1, FOXN3, and GADD45B) in 1 SARS-CoV-2 transcriptomic dataset (Fig. 3e). additionally, Fig. 3e shows the levels of downregulation of 12 genes that overlapped with vitamin D-modulated genes in cells of patients infected with SARS-CoV-2. Network analysis of these 12 genes (shown by green circles, Fig. 3f, right) indicated that several of such downregulated genes, such as ARRB1, RGS3, GGA3, HEXA, EIF3D, and PSMD3, are functionally related to the immune system (Functional enrichment analysis of network genes using Reactome with a rank of 56, p-value 1.31e-21, Supplementary Table 4). Significance of downregulation of these shared 12 genes is judged by the nature of their functions. For example, functions associated with the following genes are expected to be compromised in SARS-CoV-2-infected lung cells: (i) EIF3D, a component of the protein translational complex; reduction in the levels of this component inhibits CD8^+^ T cell activity and promotes HIV progression^38^, (ii) NLRP1, a component of inflammasome with role in innate immunity^39^, and (iii) RGS3, a G protein signaling component with role in T-lymphocyte motility during T helper 2 (Th2)-driven inflammation in airway cells^40^.

Among the 43 differentially expressed genes, 19 genes (SLPI, CCL2, STEAP4, FGL2, MARCKS, TMEM71, MXD1, DAPP1, IFITM2, OAS1, SELL, CSTA, CIR1, TIMP1, DYNLT1, IRF2, YWHAE, SCLT1, and NOP10) were upregulated in 3 SARS-CoV-2 datasets; 3 genes (GADD45B, FOXN3, and SECTM1) were upregulated in 2 datasets, and 10 genes (HCFC1R1, IER2, SIRPB1, CD4, CTSC, LIPA, PDE4DIP, ITSN1, and SSH2) were upregulated in 1 dataset (Fig. 3e). Network analysis revealed that the upregulated genes were strongly correlated with the immune system. Surprisingly, Reactome pathway enrichment analysis of network of 19 up-regulated genes identified the immune system, cytokine signaling, and adaptive and innate immunity as first four primary enriched gene sets (Fig. 3f, Supplementary Table 5). We found that many of the differentially expressed genes (that overlapped with vitamin D-modulated genes) in the cells of patients infected with SARS-CoV-2 having a role in antiviral responses were induced during viral infection. Interestingly, a subset of differentially expressed 19 genes in lung cells of patients infected with SARS-CoV-2 included genes that have been implicated in the action of interferons, development of antiviral response, and regulation of innate immune responses in individuals and cells infected with viruses^41–47^. Examples of such upregulated genes included 2’-5’-oligoadenylate synthetase 1 (OAS1), an interferon-responsive gene^48–50^, interferon regulatory factor 2 (IRF2), interferon-induced transmembrane protein 2 (IFITM2), secretory leukocyte peptidase inhibitor (SLP1), max dimerization protein 1 (MXD1), tissue inhibitor of metalloproteinase-1 (TIMP1), PH domain-containing adaptor Bam32/DAPP1, and T helper type 1 (Th1)/monocyte-associated chemokine CCL2, which have been shown to also participate in SARS-CoV-2–AEC2 signaling in lung cells^51^. In fact, upregulation of OAS1 and IFITM2 has been recently noticed as a common feature in patients infected with SARS-CoV-2^41^, and certain variants of OAS1 have been predicted to influence SARS-CoV-2 infection^52^. These observations suggest that the noticed reduced expression of the vitamin D pathway in lung cells of patients infected with SARS-CoV-2 could be associated with both down- and upregulation of cellular genes because vitamin D is known to regulate gene expression both positively and negatively. These observations suggested codysregulation of the vitamin D pathway with molecules with roles in Th1 response and immune regulation pathways in a subset of patients infected with SARS-CoV-2.

### Exploring a potential relationship between SARS-CoV-2 and PAKs

The PAK family of cellular kinases has been found to be activated by pathogens, including RNA viruses such as influenza and HIV^53,54^. Over the years, PAKs have been implicated in various stages of virus entry and replication and in supporting the cell survival phenotype during certain viral infections^20^. To understand the significance of the PAK pathway in SARS-CoV-2 pathobiology, we first examined the status of PAKs in SPPD. We found that the levels of PAK1 and PAK4 mRNAs are generally upregulated in model systems pertaining to middle east respiratory syndrome coronavirus (MERSCOV), whereas PAK2 and PAK3 are downregulated in MERSCOV models (Fig. 4a).

**Figure 4:**
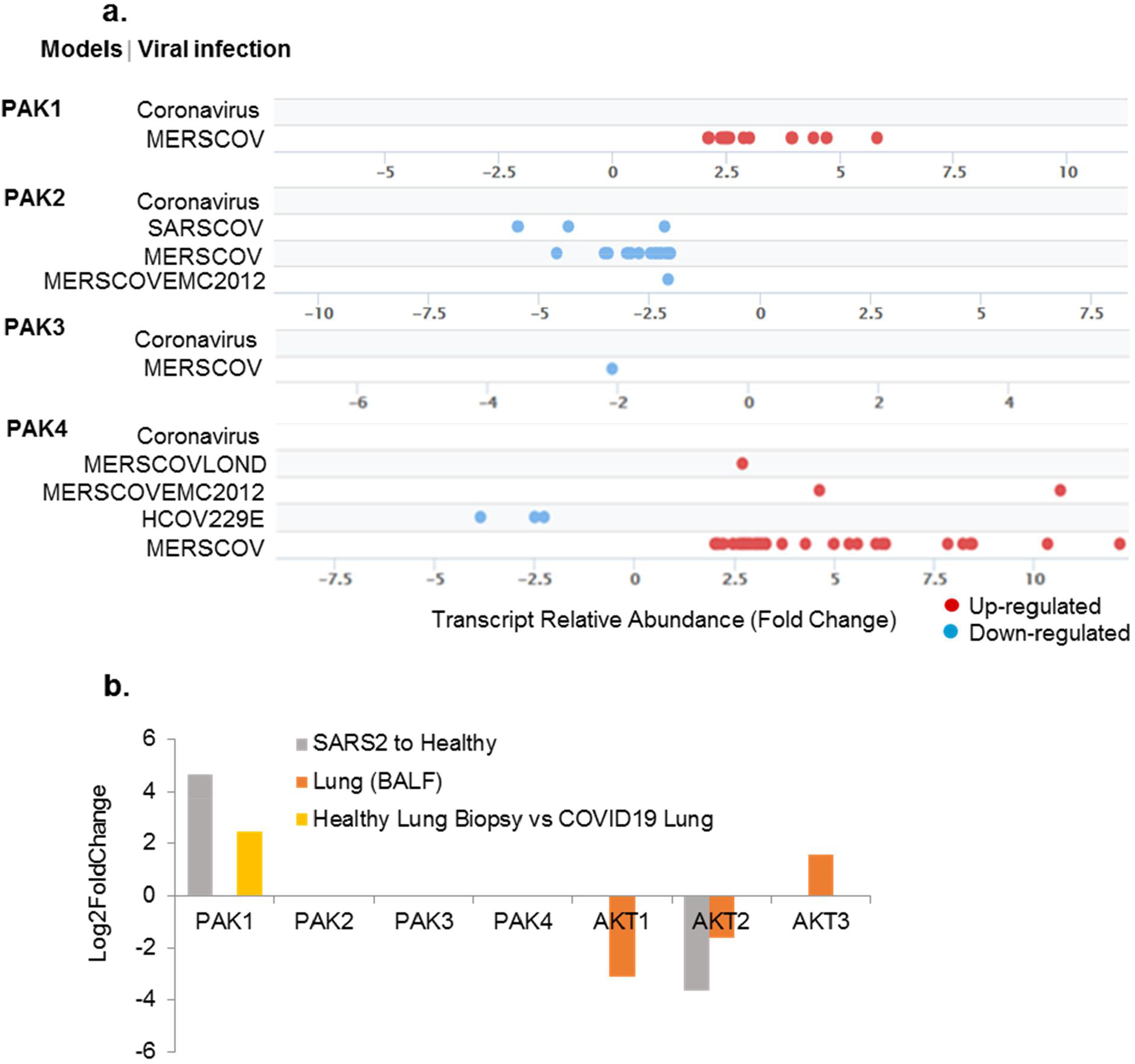
Expression of PAKs in viral infection models. **a)** Shows the fold change distribution of p21 activated kinases with an FDR cut-off of 5E-02 from viral infection from Signaling Pathway Project. **b)** Shows the expression levels of kinases in SARS-COV-2 patient samples

Further, we examined the levels PAK1–4 in transcriptomic analyses in lung cells of patients infected with SARS-CoV-2. We noticed PAK1 mRNA upregulation in datasets of all three patients infected with SARS-CoV-2, whereas PAK2 upregulation was observed in one of the transcriptomic datasets. No significant alteration was observed in the differential expression of PAK3 and PAK4 in samples collected from patients infected with SARS-CoV-2 compared with the healthy controls (Fig. 4b). In addition to PAK kinases, cell survival is known to be profoundly regulated by AKT kinases^55,56^. However, as opposed to PAKs, we noticed a reduction in AKT levels in SARS-CoV-2-infected cells (Fig. 4b). These observations suggested an association between the levels of PAK1 mRNA and SARS-CoV-2 infection. However, the current literature provides no clues about the role of PAK expression in the pathobiology of SARS-CoV-2 infection.

### A subset of differentially expressed genes in SARS-CoV-2-infected lung cells is a shared target of PAK- and vitamin D-modulated genes

Our finding of a compromised expression of several components of the vitamin D pathway and increased PAK1 expression in the same transcriptomic analyses of lung cells of patients infected with SARS-CoV-2 raised a possibility of an association between these two phenomena. We attempted to explore this possibility. In this context, previous studies have suggested a role of PAK-dependent modulation of actin polymerization in the regulation of FGF23^26,27^, a modifier of CYP27B1 and CYP24A1 in the vitamin D pathway, and the vitamin D pathway have been suggested to inhibit the levels of RAC (a positive regulator of PAK1 activation) in endometrial cancer cells^27^. A recent review article^57^ has speculated the possibility of PAK1 regulation of CCL2, a molecule shown to connect ACE2 with liver fibrosis in a murine model^58^, during SARS-CoV-2 infection. However, data supporting a connection between PAK1 and SARS-CoV-2 pathways remained uninvestigated until this point.

Because we observed a compromised vitamin D pathway and increased PAK1 expression in the SARS-CoV-2 transcriptome, we further searched for genes that might be common effectors of vitamin D and PAK pathways. Fig. 5a shows that indeed there is a widespread overlap of genes common to vitamin D- and PAK-regulated genes. Our analysis identified 563 differentially expressed genes common to vitamin D- and PAK-regulated datasets (Fig. 5b). The expression levels of the two sets of genes are depicted in the heatmap (Fig. 5c). We next searched 563 differentially expressed genes for vitamin D-modulated genes that might be downregulated or upregulated in SARS-CoV-2 transcriptome, presumably because of a compromised vitamin D pathway or gene expression due to derepression. We believe that such genes might be the shared effectors of vitamin D and PAK pathways in lung cells of patients infected with SARS-CoV-2, and hence, they might be involved in SARS-CoV-2 pathobiology.

**Figure 5:**
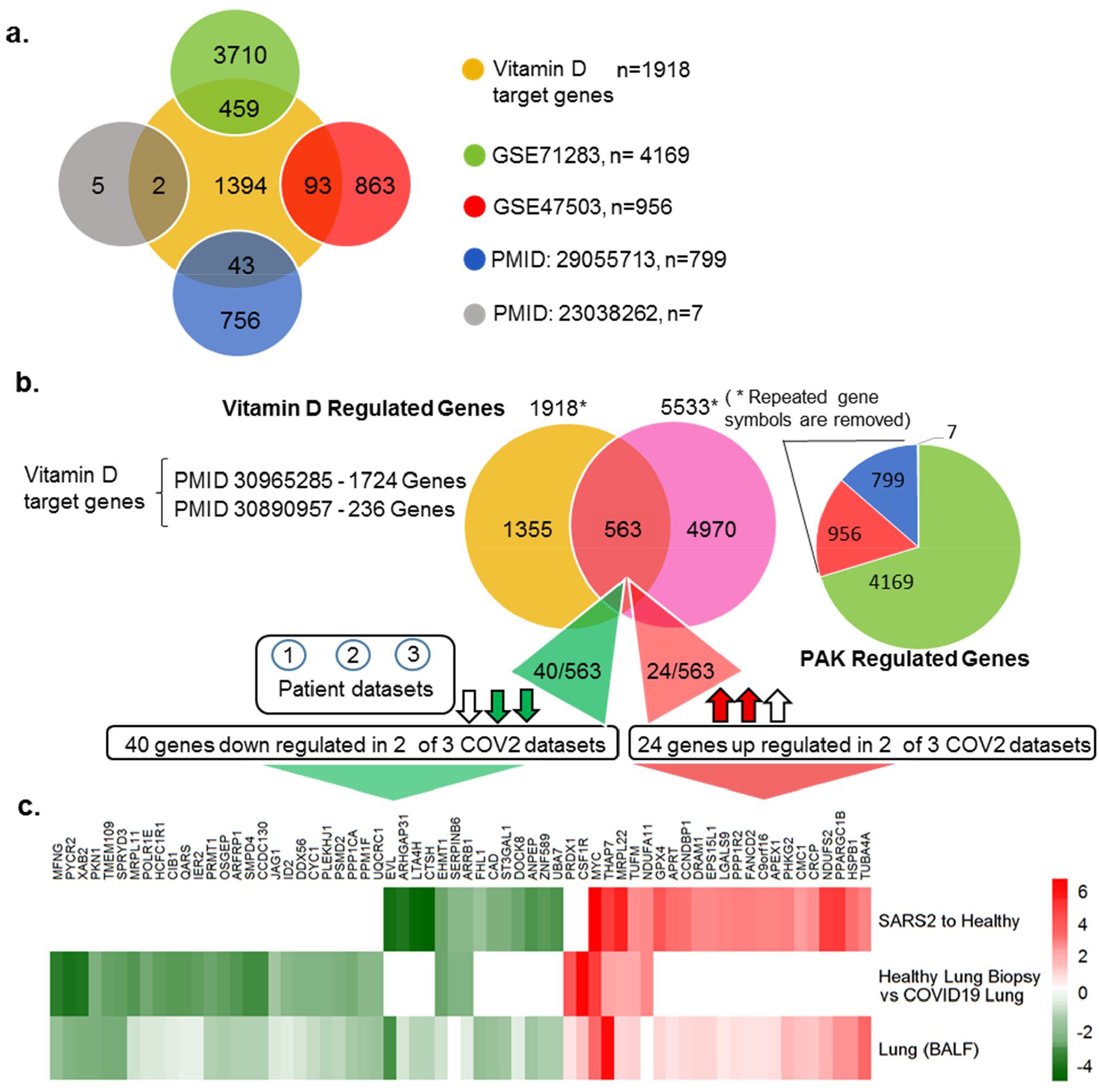
Shows the overlap between Vitamin D regulated genes and PAK regulated genes. **a)** Venn diagram shows the comparison of Vitamin D target genes for each PAK datasets and the overlapped genes are denoted using venn diagram. **b)** Venn diagram shows the overlap between Vitamin D regulated genes and PAK regulated genes. Vitamin D regulated genes are selected from RNA-Seq data from the given PubMed identification numbers (Ids) and also adopted from PANTHER database. Statistically significant genes which shown positive correlation up on PAK1 silencing and up on PAK4 over expression from given studies are considered as PAK regulated genes and a pie chart shows the number of genes selected from each PAK studies. **c)** The heat map shows the expression of 40 genes down regulated in 2 out of 3 SARS-COV-2 patient samples and the 24 genes found down regulated in 2 out of 3 SARS-COV-2 patient samples.

We focused on vitamin D-modulated genes that were downregulated in two out of three SARS-CoV-2 transcriptomic studies. Our analysis identified 40 such downregulated genes out of 563 differentially expressed genes, shared between vitamin D and PAK pathways, in SARS-CoV-2-infected lung cells (Fig. 5b). Similarly, there are 24 upregulated genes (shared between vitamin D and PAK pathways) in SARS-CoV-2-infected lung cells (Fig. 5b). These observations suggest that the evident dysregulated vitamin D pathway in SARS-CoV-2-infected lung cells could be accompanied by both down- and upregulation of downstream effectors that are also shared effectors of the PAK1 pathway in SARS-CoV-2-infected lung cells. Because SARS-CoV-2 induces PAK1 expression in lung cells and PAK1 signaling stimulates as well as represses several cellular genes^21-25^, a subset of genes in SARS-CoV-2 infected cells might be coinfluenced by a defective vitamin D and viral infection-associated PAK1 upregulation.

### Vitamin D pathway and SARS-CoV-2 pathogenesis

Aforementioned analysis identified 43 vitamin D-modulated genes dysregulated in lung cells of patients infected with SARS-CoV-2. Network analysis of 43 genes revealed that many of these genes are functionally related to NF-κB pathway and cytokine signaling, DNA damage checkpoint and response (rank 2 and 3), and apoptosis (rank 4) (Fig. 6a). Similarly, network analysis of 40 downregulated genes, which were shared targets of vitamin D and PAK pathways, in SARS-CoV-2-infected lung cells indicated that genes in the top four rank orders belong to different aspects of regulation of immune response (Fig. 6b). In contrast to the aforementioned downregulated genes, network analysis of 24 upregulated genes, which were shared targets of vitamin D and PAK pathways, suggested cell cycle regulation and gene expression as the top functional targets (Fig. 6c). These observations raised the possibility that the evident downregulation of vitamin D and upregulation of PAK1 in SARS-CoV-2-infected cells is likely to have a significant influence upon NF-κB pathway, cytokine signalling and immune regulation. The critical role of genes important for promoting the G1 -S progression as well as key components of the cell survival that could impact the pathobiology of SARS-CoV-2 is also evident in the analysis. Further studies are warranted to delineate the mechanism underlying these findings with a broad implication in the pathobiology of SARS-CoV-2.

**Figure 6:**
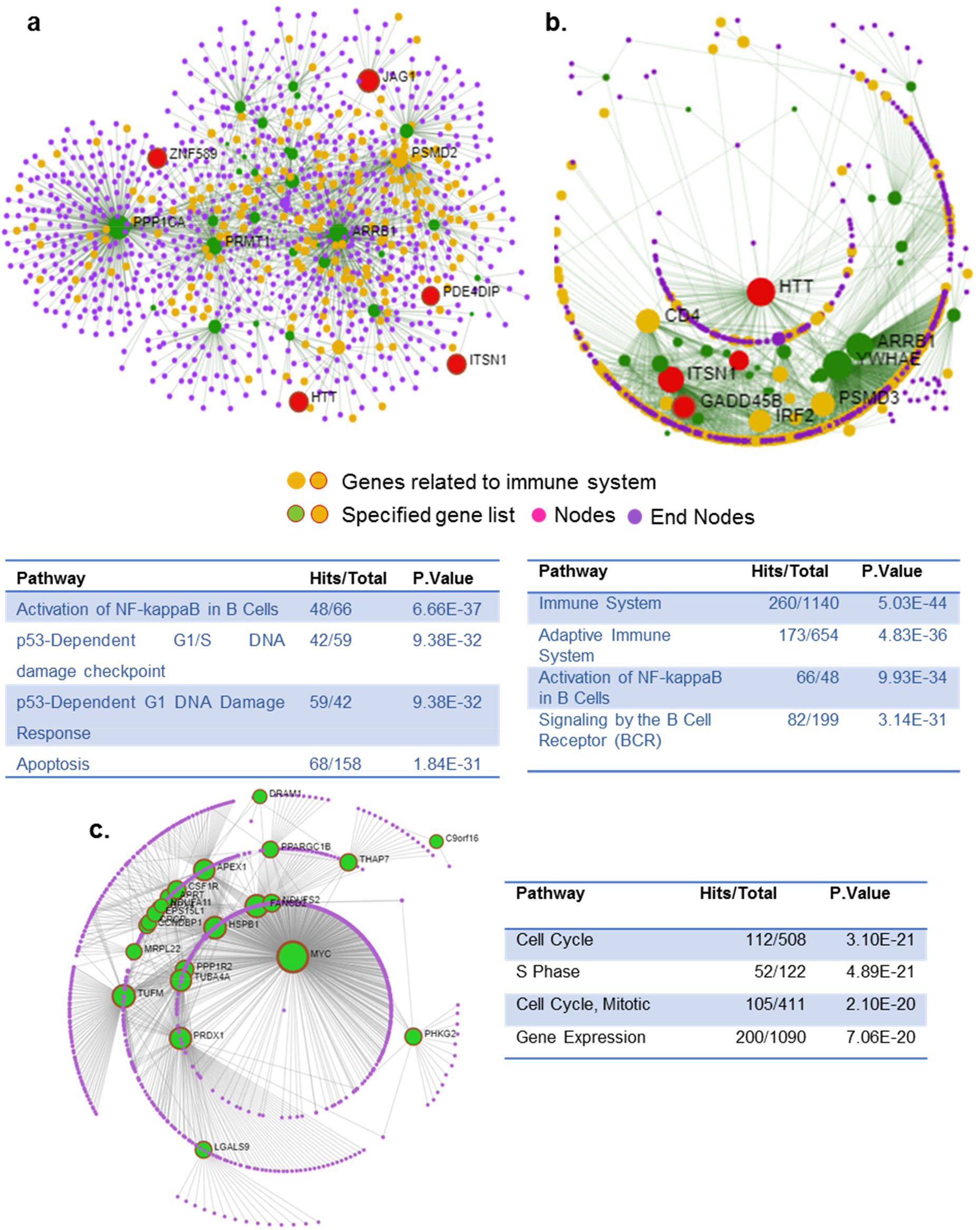
Network analysis of vitamin D-modulated differentially expressed gene in SARS-CoV-2 transcriptome. **a)** Network analysis using Network analyst tool of 40 genes shared with Vitamin D and PAK regulated genes which are down regulated in at least two SARS-COV-2 datasets. **b)** Network analysis of 43 genes shared among all 3 SARS-COV-2 datasets and Vitamin D regulated genes and the genes belong to Immune system is remarked (rank 1). **c)** Network analysis of 24 genes shared with Vitamin D and PAK regulated genes which are up-regulated in at least two SARS-COV-2 datasets. *Results of functional enrichment analysis using Reactome is provided in the table. Pathways for the first four ranks are listed with each network.

### Targets of the dysregulated vitamin D pathway with or without VDR in SARS-CoV-2-infected lung cells

The vitamin D pathway exerts its cellular effects through nongenomic/cytoplasmic and nuclear effects. Vitamin D regulates the expression of cellular genes either through modifying the expression of target genes containing vitamin D responsive elements (VDR) or indirect mechanisms as a consequence of other effects of vitamin D. VDR as a heterodimer protein complex with receptor RXR binding to the DR3-response element consisting of A/GGG/TTC/GA motif^34,59^. We next searched for the presence of VDR motifs within −1000bp and −100bp from TSS in target gene promoters among 43 differentially expressed vitamin D-modulated genes and 40 differentially expressed genes between vitamin D modulated and PAK pathways in the SARS-CoV-2-infected transcriptome. Surprisingly, we found the presence of VDR motifs in 3 of the 43 differentially expressed genes (JAG1, ST3GAL1, and ZNF589) and in 5 of the 40 differentially expressed genes (ITSN1, GADD45B, PDE4DIP, HTT, and MARCKS) in SARS-CoV-2-infected cells (Figs. 7a,b). These observations suggest that a defective vitamin D pathway could impact the levels of differential expression of vitamin D-modulated genes in SARS-CoV-2-infected lung cells via both direct mechanisms involving the VDR motif in the target genes and indirect mechanisms.

**Figure 7:**
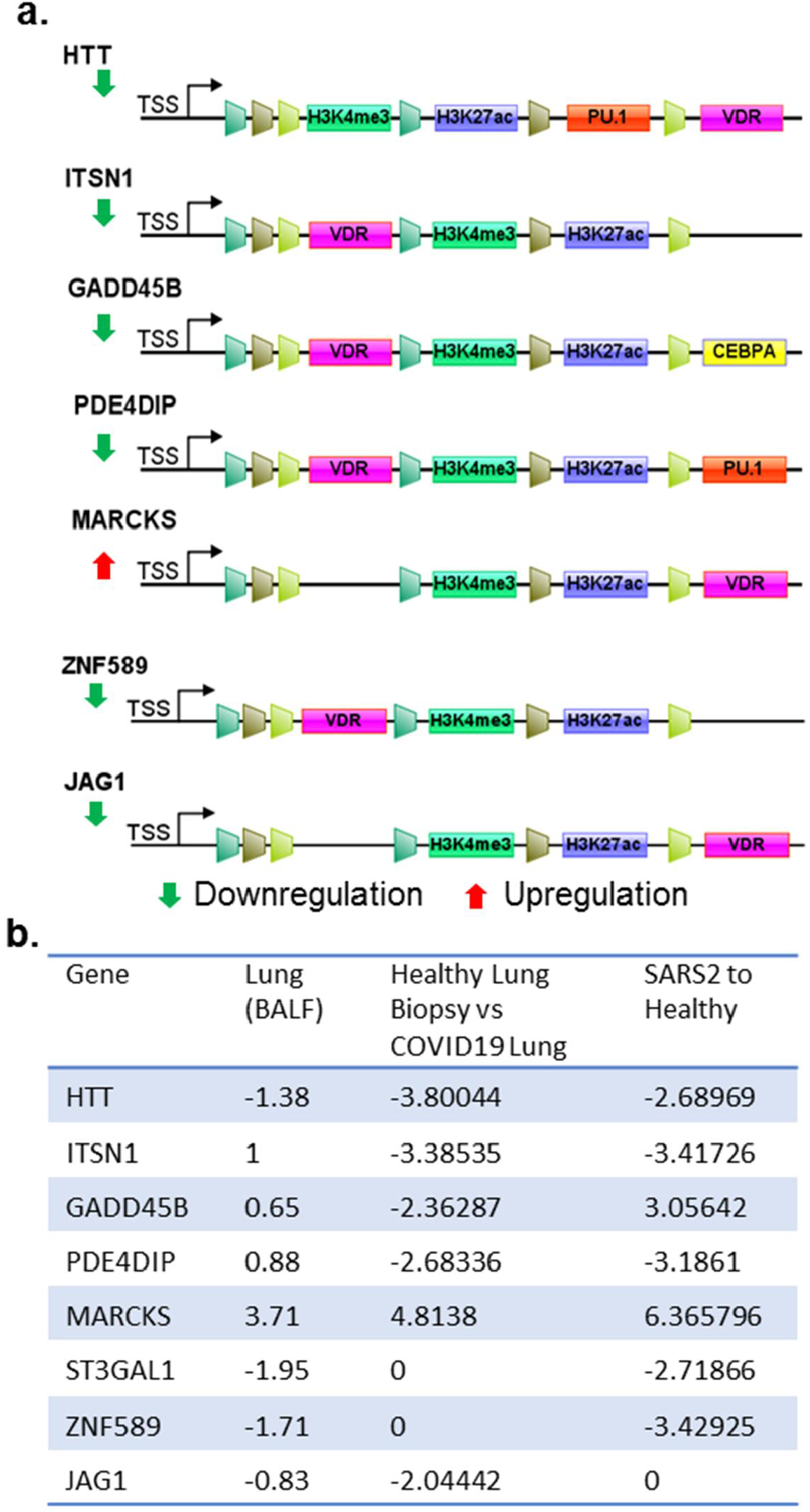
Analysis of postulated vitamin D target genes. **a)** Status of VDR binding motifs, shared histone marks and TFs in TSS as well as enhancer regions. **a)** Results when the analysis carried out for genes with VDR binding motifs. **b)** Expression level of the genes in SARS-CoV-2 patient datasets are provided.

## Discussion

In brief, the present study was conducted to assess the status of the vitamin D pathway in patients infected with SARS-CoV-2 using public transcriptomic datasets and computational approaches. Results presented support the notion of a potential association between SARS-CoV-2 infection and reduced expression of several components of the vitamin D pathway. As expected, the dysregulated vitamin D pathway in the lung cells of patients infected with SARS-CoV-2 was accompanied by dysregulation (both down- and upregulation) of cellular genes of vitamin D-modulated genes, including molecules involved in Th1 response and immune regulation. Another novel notable observation is the upregulation of PAK1 expression (but not another family of survival AKT kinases) in the same SARS-CoV-2 sample sets with a reduced expression of vitamin D pathway components, suggesting a potential correlative relationship between these two phenomena. Consistent with this hypothesis, we noticed that indeed a subset of differentially expressed genes in lung cells of patients infected with SARS-CoV-2 is also the shared target of PAK- and vitamin D-modulated genes and such genes were coinfluenced by not only a defective vitamin D pathway but also PAK1 upregulation associated with SARS-CoV-2-infection; thus, they are involved in SARS-CoV-2 pathobiology. Future studies are required to define the precise levels of crosstalk between PAK1 and vitamin D pathways and to determine whether PAK1 plays a role in the dysregulated vitamin D pathway.

As vitamin D modulates the expression of several cellular genes through a direct mechanism involving VDR motifs in the putative target genes as well as through indirect mechanisms, we observed that most differentially expressed genes, common to vitamin D and/or PAK-modulated genes, in SARS-CoV-2-infected lung cells lacked the VDR motif in target gene promoters. Hence, they were expected to be regulated by indirect mechanisms in SARS-CoV-2-infected cells by vitamin D. However, our analysis did discover, at least, eight new vitamin D target genes with a conserved VDR motif within −1000 and −100 from TSS in gene promoters as differentially expressed genes in SARS-CoV-2-infected lung cells. This raised the possibility of direct mechanisms of regulation of such vitamin D-modulated genes by the dysregulated vitamin D pathway.

To highlight the potential significance of a subset of eight newly recognized VDR-containing target genes in SARS-CoV-2 pathobiology, we here briefly discuss the potential connection between six differentially expressed genes in SARS-CoV-2-infected lung cells; however, ZNF589 and HTT remained novel in the context of viral infection.

### Intersectin-1 (ITSN1)

Cytoskeleton remodeling and interactions, including dynamics of actin polymerization–depolymerization, which is a process also regulated by PAK signaling^21^, plays a role in the early steps of endocytosis during viral infection and phagocytosis^60,61^. One such regulator is the guanine nucleotide exchange factor ITSN1 with an established role in vaccinia infection-associated actin polymerization and Fc-gamma receptor-mediated phagocytosis^62^. In the context of SARS-CoV-2 infection, actin remodeling represents a central event in inflammatory responses in the lungs^63^ and might be involved in virus entry^64^. Further, components of actin remodeling have emerged as important interactors of membranous ACE2^65^, which is the primary receptor for SARS-CoV-2. Likewise, Fc-gamma-mediated phagocytosis, a process known to be regulated by ITSN1, plays an important role in immune responses during SARS-CoV-2 infection^66^. These observations in conjunction with the identification of ITSN1 as one of the vitamin D-modulated differentially expressed genes in SARS-CoV-2-infected lung cells further raise the possibility of connecting a dysregulated vitamin D pathway with misregulation of any of the aforementioned cellular functions of ITSN1.

### Growth Arrest and DNA Damage Inducible Beta (GADD45B)

GADD45B and GADD45A have been shown to be important in the regulation of DNA damage response and senescence^67^. Relevance of the presence of a conserved VDR motif in the promoter region of GADD45B and its differential expression in SARS-CoV-2-infected lung cells (as shown in this study) lies in a recent finding presenting the differential expression of GADD45B as well as other genes belonging to mitochondrial functions in patients with chronic obstructive pulmonary disease (COPD)^68^. Further, as older patients with COPD are more susceptible to SARS-CoV-2 infection^69^, GADD45B might be involved in SARS-CoV-2 pathobiology in patients with COPD.

### Phosphodiesterase 4D-interacting protein (PDE4DIP)

PDE4DIP is an understudied enzyme that hydrolyzes the 3′ cyclic phosphate linkages in 3′,5′-cAMP and cGMP and 3′,5′-cAMP and participate in cellular signaling and other processes^70^. PDE4DIP has recently been suggested to be a differentially expressed genes in lung cells in COPD^71^. As no precedence of vitamin D regulation of PDE4DIP exists, our present finding indicates the possibility of a regulatory relationship between vitamin D and cyclin AMP or GMP signaling in SARS-CoV-2 pathobiology in lung cells.

### ST3 Beta-Galactoside Alpha-2,3-Sialyltransferase 1 (ST3GAL1)

ST3GAL1 is an enzyme with sialyltransferase activity. It participates in transferring sialic acid to substrates with galactose as the acceptor and is involved in glycosylation^21^. Glycosylation is fundamental to the regulation of numerous cellular processes, including vitamin D-binding proteins^72^. As certain genetic variants of ST3GAL1 have been shown to be associated with an increased probability of severity in patients with influenza A(H1N1)pdm09 infection^73^, the dysregulated vitamin D pathway in SARS-CoV-2-infected patients might be accompanied by misglycosylation of certain proteins, which in turn may impact SARS-CoV-2 pathobiology in lung cells.

### Jagged-1 (JAG1)

Activation of Notch receptors by Notch ligands, including JAG1, and resulting signaling events have been shown to be involved in cell-to-cell communication. In addition, JAGs/Notch signaling influences diverse aspects of cytokine biology, inflammation, dendritic cell biology, T-cell development, B-cell repertoire, and innate immunity^74–77^. Further, Notch signaling has been suggested to be downstream of IL6, which is a proinflammatory cytokine with a role in the inflammatory storm, in the lung and heart^78^. In the context of viral infection, JAG1/Notch 1 signaling participates in the development of Th2 response in bronchial epithelial cells following infection with respiratory syncytial infection^79^. Interestingly, experimental downregulation of JAG1has been shown to promote Th1 response while suppressing Th2 differentiation, a situation similar to that observed in SARS-CoV-2-infected lung cells in this study^79^. This situation is also somewhat similar to that of SARS-CoV-2-specific CD4^+^ T cells and is shown to be Th1 type^80^. These observations and the finding in this study that JAG1 could be a vitamin D-regulated gene suggest that a dysregulated vitamin D pathway might influence cellular immunity via the JAG1 pathway.

### Myristoylated alanine-rich C kinase substrate (MARCKS)

MARCKS, a substrate of protein kinase C and actin-interacting protein, is present in the plasma membrane. It modulates vascular contractility and regulates Ca^2+^ and phosphatidylinositol 4,5-bisphosphate signaling^81^. Upon activation by PKC signaling, MARCKS translocates from the plasma membrane to the cytoplasm and regulates many cellular processes depending on actin remodeling, such as membrane trafficking and phagocytosis. MARCKS has been shown to be involved in lung diseases, including COPD and asthma^82^, probably because of its ability to regulate mucin secretion and inflammation^83^. Activation of PKC, either directly or through secreted cytokines, is generally viewed a common event during viral infections, including human immunodeficiency virus in promonocytic cells, respiratory syncytial virus in bronchial epithelial cells^84,85^, and hepatitis B virus transactivator HBx^86^. Additionally, Ang II has been shown to activate PKC-β, and in turn, cellular redistribution of MARCKS in neurons^87^. Vitamin D3 metabolites are known activators of PKC, and vitamin D deficiency is associated with a dysregulated subcellular distribution of PKC isoforms in rat colonocytes^88^. The aforementioned observations suggest that a defective vitamin D pathway might be associated with a misregulated subcellular distribution of MARCKS in lung cells and could be involved in SARS-CoV-2 pathobiology.

In brief, this study provides insights about a potentially causal association between a compromised vitamin D pathway at multiple levels and SARS-CoV-2 infections in patients, and consequently, dysregulation of pathways downstream to vitamin D. These preliminary results now set the stage for experimental validation of the observations and postulations made using computational approaches. Additionally, this study also sets the stage for conducting a larger study to determine whether a compromised vitamin D pathway might influence the susceptibility of lung cells to SARS-CoV-2 and/or the consequence of SARS-CoV-2 infection.

## Methods

### Gene Expression profiling in viral infection models

Relative transcript abundance for individual genes were analysed using Ominer tool from The Signaling Pathways Project^89,90^. Single gene target was analysed under transcriptomics category. An FDR cut-off of 5E-02 was applied for the analysis, expression profiles for viral infection models were extracted and represented in the article.

### SARS-CoV-2 lung patient transcriptome

High throughput transcriptomic data for SARS-COV-2 patient samples were selected from the studies submitted under Genome expression studies related to SARS-CoV-2 in GEO up to July 2020, wherein first study performed a transcriptomic profiling of bronchoalveolar lavage fluid (BALF) was performed on SARS-CoV-2 patient samples and healthy donors from Zhongnan Hospital of Wuhan University^32^. Second study reports that BALF cell transcriptome indicates robust innate immune responses in SARS-CoV-2 patients^30^ and the remaining studies provided the transcriptomic data ofSARS-CoV-2 deceased patient, the data has accessed from Gene Expression Omnibus under the accession GSE147507, GSE153970, GSE150316 and GSE152075. Gene expression data were filtered for SARS-CoV-2 lung and normal lung and sample size is restricted to 5 in the case of datasets with large samples size.

### SARS-COV-2 lung cell line transcriptome

The transcriptome data of primary human lung epithelium (NHBE), transformed lung alveolar (A549), transformed lung-derived Calu-3 cells mock treated or infected with SARS-CoV-2 (USA-WA1/2020), data was accessed from Gene Expression Omnibus under the accession GSE147507^31^.

### PAK silenced transcriptome data

The gene expression profiles of MCF10A.B2 cells (MCF10A cells expressing a chemically activatable form of Her2) stably expressing a Tet inducible shRNA directed against Pak1 gene was accessed from Gene Expression Omnibus under the accession GSE71283^91^.

### Differential Expression Analysis and Data

Differential expression data available as supplementary data were collected and differential expression analysis was performed for the gene count provided under the accession GSE147507 using BioJupies^92^.

## Functional Analysis

Gene ontology analysis as well as COVID19 enrichment analysis was performed using Metascape^37^.

## Network analysis

Network analysis for the set of genes were performed using Network Analyst 3.0 tool^93–95^ with official gene symbol for generic protein-protein interactions using IMEx Interactome^96^and default confidence score cutoff (900). Corresponding gene cluster was analysed for biological pathway involvement using Reactome database^97^.

## Acknowledgements

Bijesh George and Aswathy Mary Paul are supported by fellowships from the Rajiv Gandhi Centre for Biotechnology.

## Authors Contributions

R.K. conceived, designed and directed the study; R.K. and M.R.P. analyzed the data; R.K., M.R.P. and T.R.S.K. edited the draft manuscript. B.G. performed bulk of the computational studies; A.M.P. performed PubMed analysis; and A.R. analyzed promoters of select differentially expressed genes. All authors reviewed the manuscript.

